# Genetic Linkage Studies in F_2_ Cowpea (*Vigna Unguiculata* (L.) Walp) Mapping Populations Using Qualitative Morphological and SNP Markers

**DOI:** 10.64898/2025.12.19.695351

**Authors:** Henry K. Mensah, Ralieva A.K. Norety, Isaac K. Asante, Felicia Oppong

## Abstract

The research was conducted to study genetic linkage in 45 F_2_ cowpea mapping populations derived from a cross between two inbred lines Golinga (cultivated variety) and a Wild relative. The data used for the study were based on 13 polymorphic SNPs and 17 morphological markers. The segregation ratios were analyzed revealed that five traits (growth habit, growth pattern, leaf shape, plant hairiness and pod hairiness) segregated significantly according to 3:1 mendelian classical ratio for inheritance whilst the remaining 13 showed 9:7 ratios. Genetic linkage mapping was performed by using the software QTLiCIMapping version 4.2. All the markers used were polymorphic. Eleven linkage groups were detected which spanned a total map length of 93.8217 cM at a LOD score of 4. Variation in marker distribution on all the 11 linkage groups was observed. Fourteen markers were distributed on Chromosome 1. In all cases, segregation distortion was observed for all the markers in each of the linkage groups. Segregation distortion in Chromosome 1, Chromosome 5, Chromosome 8 and Chromosome 11 were highly skewed to the wild parental type. This work provides one of the few integrated morphological–SNP linkage maps for cowpea and highlights genomic regions associated with non-Mendelian inheritance. These findings supply foundational genetic information for QTL discovery, marker-assisted selection, and breeding strategies aimed at improving yield, nutrition, and climate resilience in cowpea.

**Author Summary:** In this study, we set out to understand how important physical traits in cowpea are passed from one generation to the next, and to build a simple genetic map that can guide future crop improvement. Cowpea is a major source of food and nutrition for millions of people, especially in Africa, yet progress in improving the crop has been slowed by limited genetic information.

By crossing a cultivated cowpea variety with its wild relative, we examined how traits such as growth form, leaf shape, pod appearance, and seed characteristics segregate in their offspring. We also combined these observations with DNA markers to create a genetic map showing how these traits and markers are arranged along the chromosomes.

Our results show that some traits follow the classic patterns of inheritance, while others are influenced by interactions between pairs of genes. We also discovered regions of the genome where inheritance was skewed toward one parent, offering clues about underlying biological processes. By providing one of the few combined morphological and DNA maps for cowpea, our work creates a foundation that breeders and researchers can use to develop improved, desired, and climate-resilient cowpea varieties.

## Introduction

Genetic mapping (also known as genetic linkage mapping or meiotic mapping) is one of the various applications of molecular markers in any species. It refers to the determination of the relative positions of genes on a DNA molecule (Chromosome or plasmid) and the distance between them [1] The genetic map indicates the position and relative genetic distances between markers along chromosomes [2]

The construction of detailed genetic maps with high levels of genome coverage is the first step for some of the applications of molecular markers in plant breeding [3] and serves five purposes: allow a detailed genetic analysis of qualitative and quantitative traits that enable localization of genes or quantitative trait loci [4]; facilitate the introgression of desirable genes or QTLs through marker-assisted selection; allow comparative mapping between different species to evaluate the similarity between genes orders and function in the expression of a phenotype [5,6]; provide a framework for anchoring with physical maps based on Chromosome translocations, DNA sequence, or other direct measures [4], constitute the first step towards positional or map-based cloning of genes responsible for economically important traits [7]. Genetic-map construction is a critically important tool for further genomic studies, as well as for genetic breeding of economically important species such as cowpea[1].

Cowpea (*Vigna unguiculata* L. Walp) is one of the most ancient crops known to man. Cowpea is also a widely grown legume in tropical and subtropical Africa where it is mostly cultivated by small-scale farmers, usually intercropped with maize or sorghum. The seeds are an excellent source of carbohydrate (50–60%) and an important source of protein (18–35%) [8] They also contain an appreciable quantity of micronutrients such as vitamin A, iron and calcium [9]. It contains significant amounts of polyphenolic compounds that are beneficial to human and animal health including phenols, flavonoids and tannins [10,11]. Cowpeas contain bioactive antioxidants such as vitamin C, carotenoids and phenolic compounds [10].

Highly saturated genetic linkage maps are extremely helpful to breeders and are an essential prerequisite for many biological applications such as the identification of marker-trait associations, mapping Quantitative Trait Loci (QTL), candidate gene identification, development of molecular markers for Marker-Assisted Selection (MAS) and comparative genetic studies [1]. With the rapid development of biotechnology, dominant DNA markers were gradually replaced by co-dominant markers in genetic mapping including Single Nucleotide Polymorphisms (SNPs) and simple-sequence repeats (SSRs) OR microsatellites. Thus, this research work aimed at constructing a genetic linkage map of cowpea using morphological and SNP makers to facilitate the development of improved cowpea varieties to ensure food security and combat malnutrition globally.

## Results and Discussion

### Segregation pattern of makers

Five traits, namely: growth habit [*GH*], growth pattern [*GP*], leaf shape [*LSH*], plant hairiness [PLH] and pod hairiness [*PDHN]* segregated significantly according to 3:1 mendelian classical ratio for inheritance whilst the remaining 13 showed 9:7 ratios (Table 1). Chi-square segregation was calculated at a *P-value* of 0.05 and one and two degrees of freedom. Since all the recorded chi-square values were below that of the expected value, the null hypothesis was failed to be rejected *i*.*e*., there were significance difference between the observed and expected ratios.

**Table 1:** Segregation pattern for qualitative markers.

| Marker | Trait | Ratio | $\chi^2$ |
| --- | --- | --- | --- |
| *GH* | Growth habit | 3:1 | 0.1773* |
| *GP* | Growth pattern | 3:1 | 3.1277* |
| *TT* | Twinning tendency | 9:7 | 0.5677* |
| *PP* | Plant pigmentation | 9:7 | 0.0165* |
| *LSH* | Leaf shape | 3:1 | 3.1277* |
| *LSZ* | Leaf size | 9:7 | 0.5677* |
| *PLH* | Plant hairiness | 3:1 | 2.0496* |
| *LC* | Leaf colour | 9:7 | 0.5137* |
| *LT* | Leaf texture | 9:7 | 0.0274* |
| *LM* | Leaf markings | 9:7 | 2.5562* |
| *PTH* | Petiole thickness | 9:7 | 0.5137* |
| *RP* | Raceme position | 9:7 | 0.5137* |
| *PDAT* | Pod attachment | 9:7 | 0.5137* |
| *PDC* | Pod curvature | 9:7 | 0.5137* |
| *PDTH* | Pod thickness | 9:7 | 0.5137* |
| *PDCL* | Pod colour | 9:7 | 0.5677* |
| *PDHN* | Pod hairiness | 3:1 | 2.0496* |
| *PTC* | Pod tip colour | 9:7 | 0.0274* |

\*= significant at 0.05 or 5% level

The results of the current study support monogenic control for growth habit, growth pattern, leaf shape, plant hairiness and pod hairiness. [12] working on cowpea pod shape reported 9:7 ratios for straight pod to curved pods. This agrees with the results obtained in the current study where F_2_ ratio of 9:7 was obtained for straight pods to curved pods. This signifies the presence of epistatic interaction of duplicate recessive genes for straight-shaped pods.

[13] also reported digenic control for pod colour in cowpea with a segregation ratio of 9:7 implying the presence of epistatic gene interaction. Similarly, this present study observed a ratio 9:7 ratio in F2 progenies. This shows the dominance of pigmentation over non-pigmentation, which is in agreement with the work of [14], who reported that wholly purple pods are dominant over green pods.

Furthermore, in the study of unripped pod tip pigmentation by [13], the F2 segregated 9:7 ratios. This corroborates with the finding of this study where pod tip colour the F_2_ segregated fitted closely to a 9:7 ratio. [13] noted that two pairs of genes appeared to control the trait in a complementary manner.

### Construction of the genetic linkage map

Thirteen polymorphic SNP markers and eighteen main morphological markers were used to generate eleven (11) linkage groups at a LOD score of 4.0 (Figure 1). The map contained 31 markers spanning a Chromosome length of 93.82 cM at an average marker length of 3.13 cM (Table 2). Chromosome 1 contained 14 markers spanning a Chromosome length of 57.94 cM at an average marker length of 4.13 cM Six of the eleven Chromosomes had 1 marker each. Chromosomes 4 and 7 had two markers each spanning total length of 25.19 cM and 10.69 cM, respectively. The long arm of Chromosome 1 had 10 markers while the short arm had 4 markers (Figure 1).

**Table 2:**
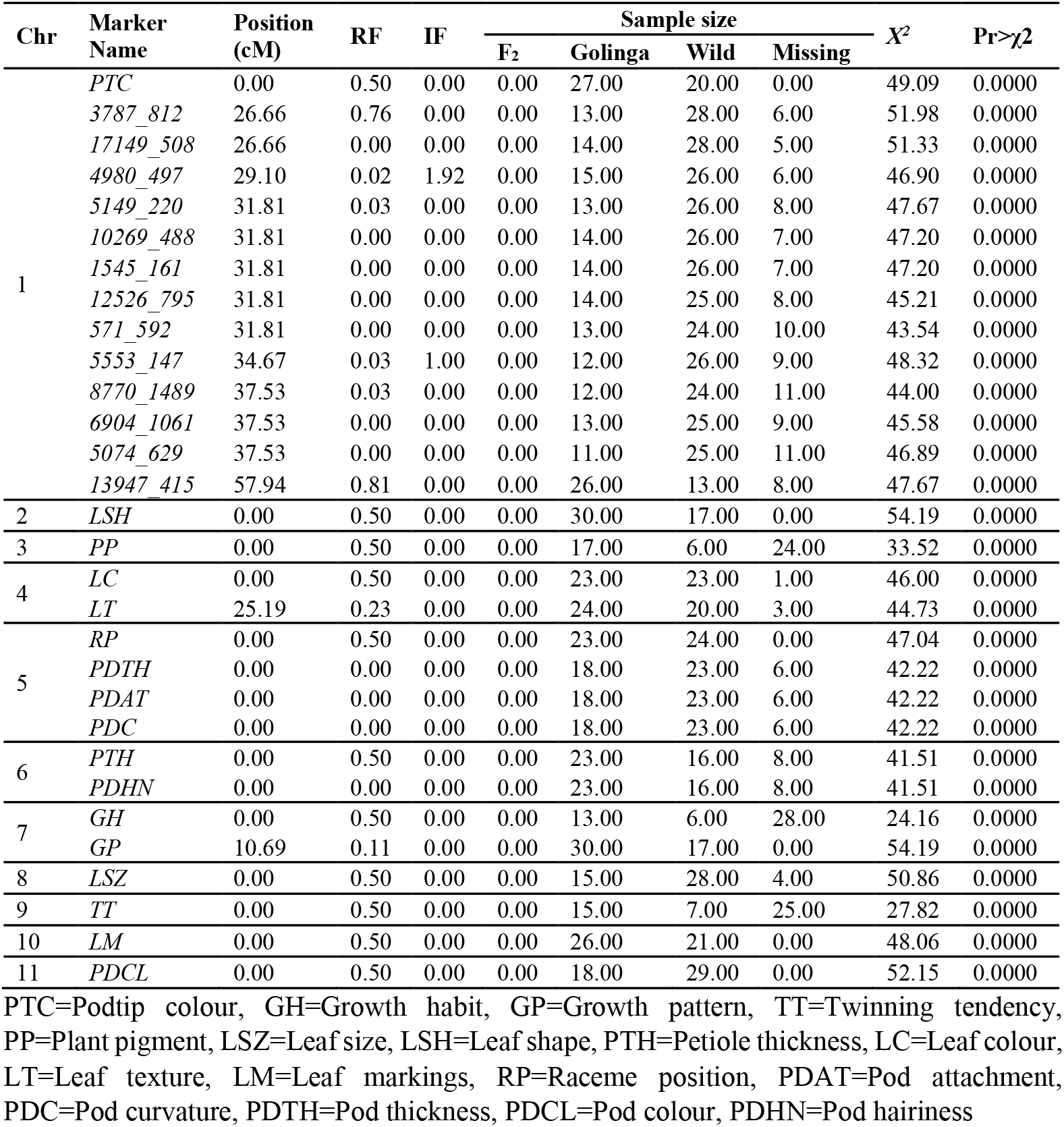
Marker information and test of segregation distortion for morphological and SNPs markers.

**Figure 1:**
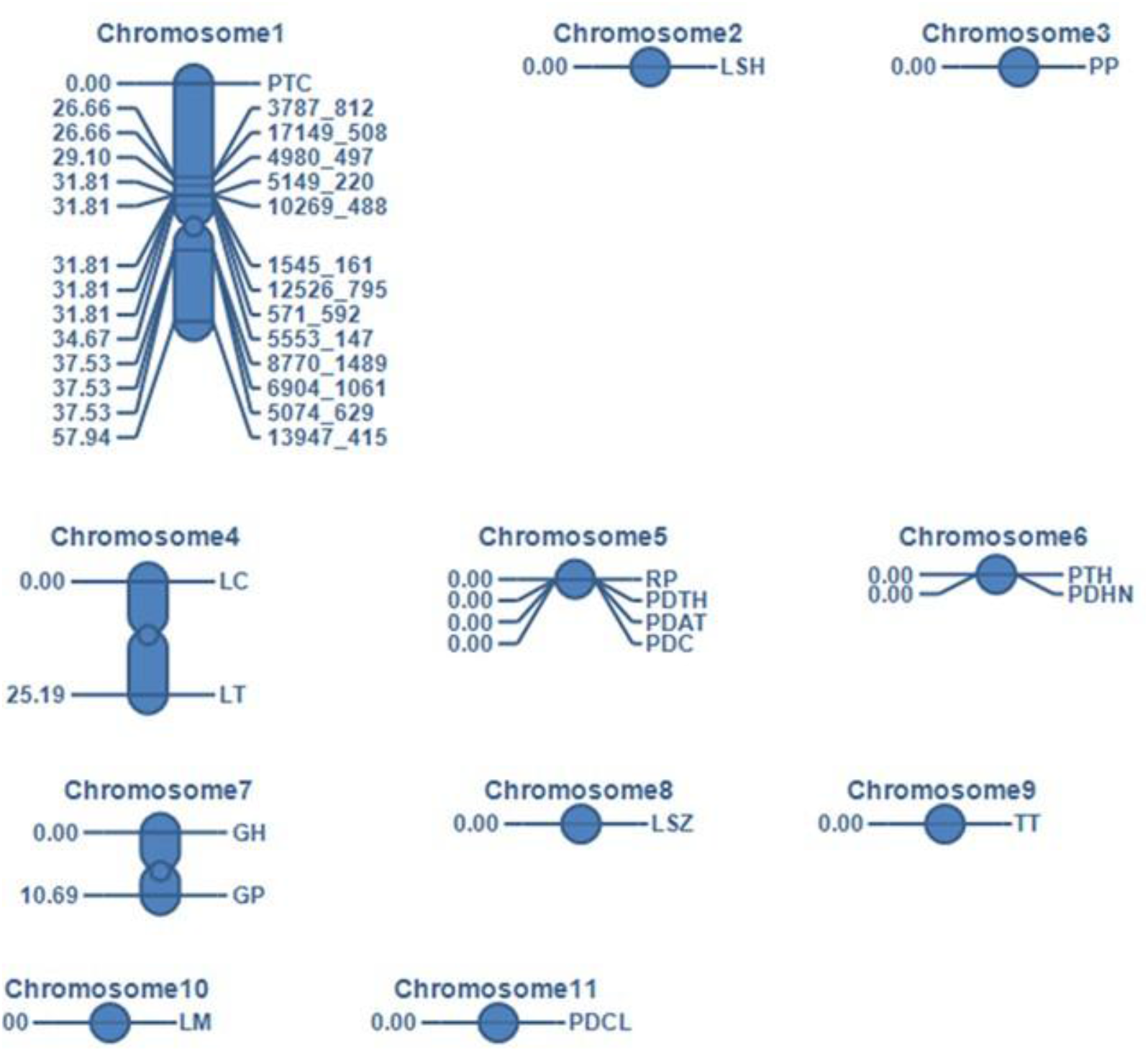
Linkage groups and their associated main morphological markers and SNPs markers. Key: PTC=Pod tip colour, GH=Growth habit, GP=Growth pattern, TT=Twinning tendency, PP=Plant pigment, LSZ=Leaf size, LSH=Leaf shape, PTH=Petiole thickness, LC=Leaf colour, LT=Leaf texture, LM=Leaf markings, RP=Raceme position, PDAT=Pod attachment, PDC=Pod curvature, PDTH=Pod thickness, PDCL=Pod colour, PDHN=Pod hairiness

The eleven linkage groups generated in the present study is at variance with the results of; [15] using 125 polymorphic SNP markers generated 12 linkage groups at a LOD score of 3.5. The findings of the current study were in disagreement with [16] who developed the first genetic map using a mapping population of 58 F_2_ plants derived from a biparental cross generated 10 linkage groups which spanned 680 cM of the cowpea genome. This present study diverged with that of [17] who generated the second cowpea genetic linkage map using 94 F8 recombinant inbred lines (RILs) derived from a cross between two cultivated genotypes *IT84S-2049* and *524B*. A total of 181 loci, comprising 133 RAPDs, 19 RFLPs, 25 AFLPs and three each of morphological and biochemical markers were assigned to 12 LGs spanning 972 cM with an average distance of 6.4 cM between markers. This second map was improved with the addition of 242 new AFLP markers generated 11 LGs spanning a total of 2670 cM, with an average distance of 6.43 cM between markers [18] similar to the number LGs formed in the current study. Comparable results were observed in the study by [19], who used AFLP and 5 cowpeas derived microsatellites to construct a linkage map using a set of RILs to generate 11 linkage groups with an average distance between markers of 9.6 cM.

These variances in reports may be due to the sample size, differences in genotypes, the differences in marker types and the number of markers used.

In addition, maps developed from crosses between cultivars are most useful for breeding applications as they identify markers that are polymorphic within the cultivated gene pool and are therefore more likely to be present in other cultivated X cultivated crosses used by breeders.

### Segregation distortion linkage

There was significant segregation distortion (P=0.0000) at all the thirty gene loci. Segregation distortion in Chromosome 1, Chromosome 5, Chromosome 8 and Chromosome 11 were highly skewed to the wild parental type whereas Chromosome 2, Chromosome 3, Chromosome 6 and Chromosome 10 skewed to the Golinga parental type (Table 2).

In a study by [20], they constructed genetic linkage map and conducted genetic analysis of domestication related traits in mung bean noted segregation distortion (P=0.05) was observed for 61 markers. Most of the markers were on Linkage group (LG) 4 showed a high level of segregation distortion (P=0.0000) toward a higher-than-expected frequency of the Wild-parent allele. In contrast with this study which reported segregation distortion (P=0.0000) was observed in 30 markers with most of the markers observed on Chromosome 1 towards a lower than expected frequency of the wild-parent allele which suggests the association of these loci in the preferential transmission of the paternal line. Deviation from normal assortment may be due to linkage of some traits and this type of distorted ratios have been observed in studies by [21] in chickpea; [22] in *Phaseolus* and [23] in blackgram.

All the markers located on Chromosome 1, Chromosome 5, Chromosome 8 and Chromosome 11 exhibited strongly distorted segregation with respect to the expected Mendelian inheritance, towards the male parental line (wild). This skewed segregation was also observed in a second F_2_ population of 45 plants derived from the biparental cross, confirming the presence of a region of distorted segregation on this linkage group and its heritability. The most skewed markers which were tightly linked to each other, indicated that they may both be closely associated with the genetic factor responsible for segregation distortion in cowpea. According to [24,25], the distorted segregation could be caused by gametophytic factors that affect either male or female gametes. Pollen fertility and meiosis can be analysed to determine their relationship with segregation distortion.

## Materials And Methods

### Development of mapping population

Forty - five F_2_ mapping populations (accR1-accR45) of cowpea were developed from a biparental cross between Golinga (accR46) and a Wild relative (accR47). These accessions were obtained from the Genetics Lab in the Department of Plant and Environmental Biology (DPEB). A single row plot design was used. Two seeds from each F_2_ population and the two parental checks (Golinga and Wild) were planted in a pot. A distance of 1m x 1m was maintained between the pots. The 5-row plots consisted of 10 pots.

### Segregation pattern of morphological markers

Chi-square test of significance was used to study gene interactions for the F_2_ cowpea mapping populations. The segregation ratios were analyzed using the formula:

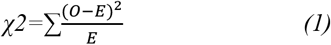

χ ^2^= Test statistic, ∑=the sum of, *O*=Observed frequency, *E*= Expected frequency

### Genotyping with single nucleotide polymorphism (SNPs) markers

Four leaf discs of 6 mm diameter were punched from 15-day old leaves and collected into 96 deep well PCR plates. Before sampling, leaves were dried at 50^°^C for 2 days to prevent possible mould development during shipment to the outsourced laboratory. DNA extraction and genotyping were done by sequencing in the United Kingdom hosted by the Laboratory of the Government Chemist, LGC. LGC genomics used a unique proprietary in-house technology (oKtopure™ protocol) to extract the total DNA. Isolated DNA was analyzed using UV spectrophotometry to estimate both the quality and quantity of the DNA, while preliminary PCR at a serial dilution was carried out and the results produced were duplicated to identify the best dilution to run the samples. The SNP genotyping was done using KASP genotyping reactions. KASP™ is also proprietary genotyping technology of LGC TM. It consists of three components namely the sample DNA, KASP assay mix and KASP master mix. KASP master mix contains two universal (FRET) fluorescent resonance energy transfer cassettes (FAM and HEX), ROX™ passive reference dye, Taq polymerase, free nucleotides and MgCl2 in an optimized buffer solution, while the KASP assay mix is specific to the targeted SNP and consists of two competitive, allele-specific forward primers and one common reverse primer. Each forward primer incorporates an additional tail sequence that corresponds to one of two universal FRET cassettes present in the KASP master mix.

### Linkage mapping construction

Screening was done to select polymorphic morphological traits and SNPs markers within the mapping populations. Out of 43 SNPs markers that were screened thirteen showed polymorphisms (Table 3). Eighteen polymorphic morphological markers were scored (

**Table 3:**
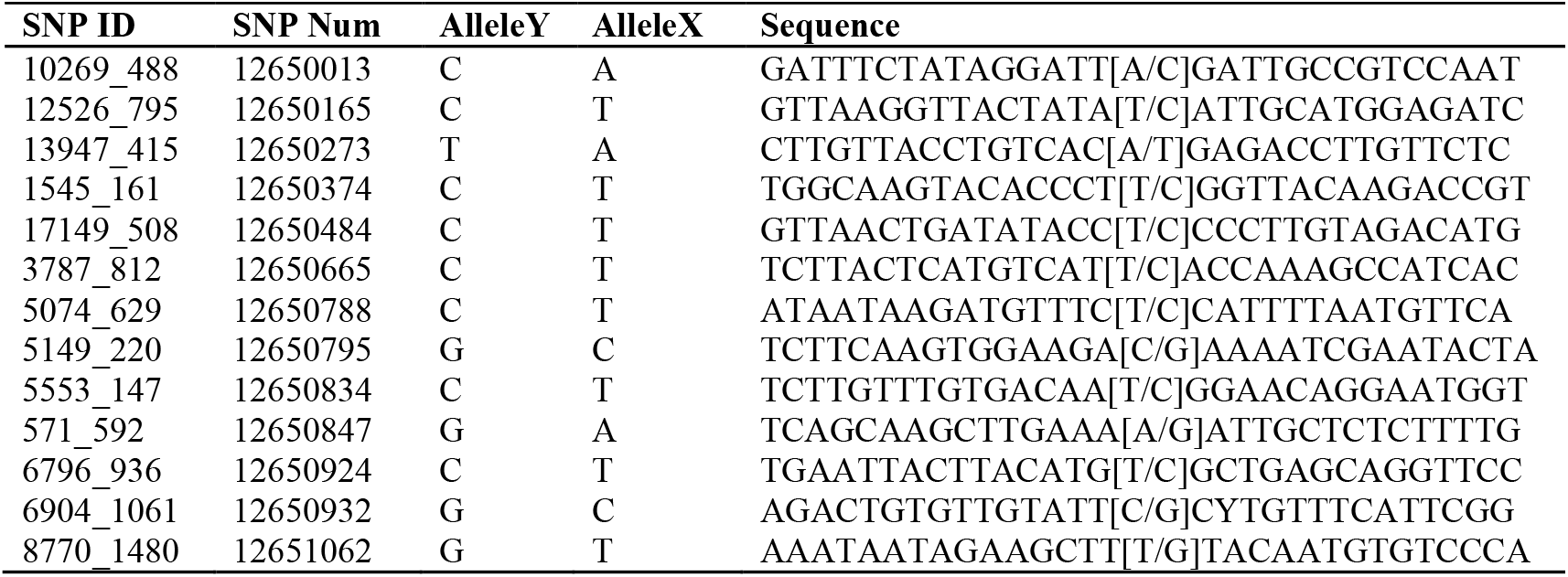
SNPs markers used for Linkage map construction.

### Segregation distortion linkage

Segregation distortion linkage was performed by using the software QTL IciMapping version 4.2. The Interval Mapping Method was used for the analysis. A segregation distortion linkage map was constructed using the eighteen morphological and the 13 SNPs markers. (**Table *4***, S1). The QTL IcMapping software version 4.2 [26] was used for the linkage map construction. The ‘Group’ command was used to identify linkage grouped and the ‘Order’ command was used to establish the most likely order within each linkage group. The orders were confirmed by permuting all adjacent markers by the ‘Ripple’ command. LOD score of 4.0 was used as the threshold for detection of linkage. The analysis was performed by using the Inclusive Composite Mapping with additive and dominance ICIM ADD method.

**Table 4:**
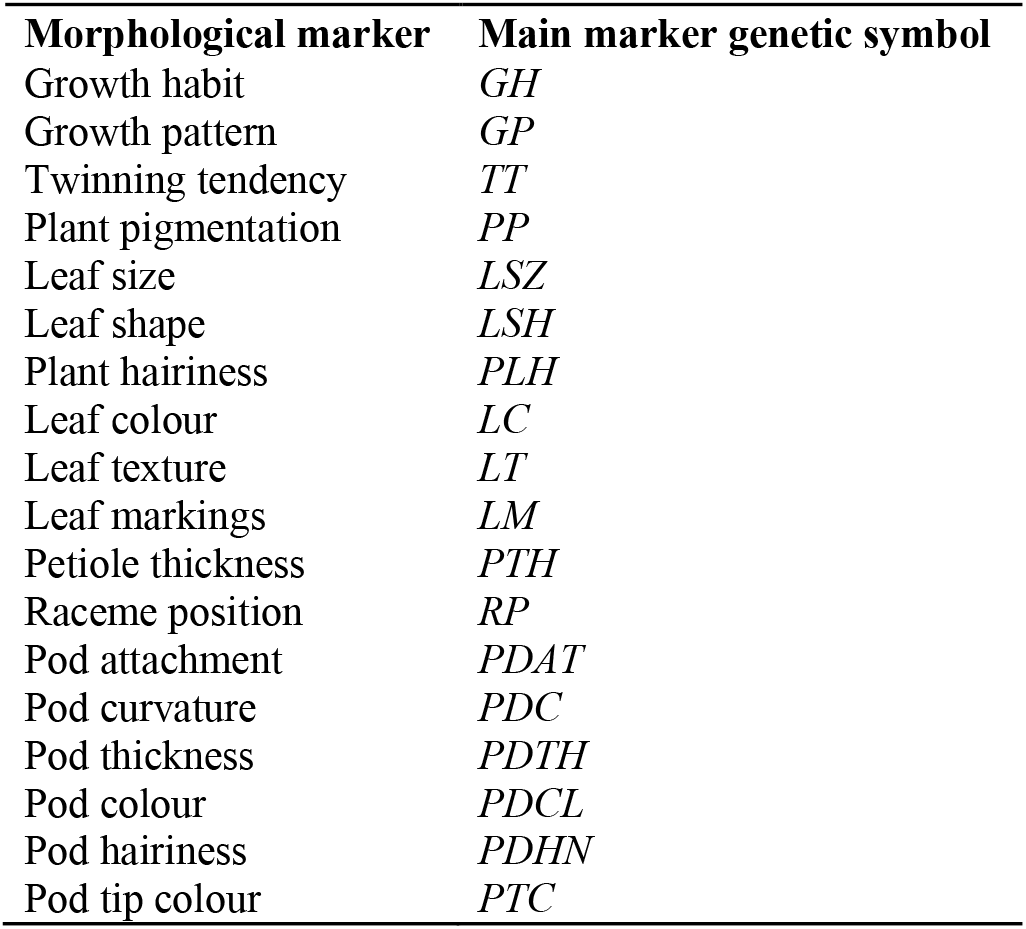
Morphological markers for linkage map construction.

## Conclusion

All morphological traits studied segregated significantly according to either 3:1 or 9:7 mendelian classical ratio for inheritance. Eleven chromosomes of cowpea (*Vigna unguiculata)* were detected which spanned a total map length of 93.8217 cM at a LOD score of 4. Markers located on Chromosomes 1, 5, 8 and 11 exhibited strongly distorted segregation towards the wild parental types which was observed in the F_2_ population. The study shows that that the set of SNPs markers used in this study provides baseline information to study cowpea genome with the aim of facilitating its improvement in the breeding programmes. Therefore, SNP markers could be used to construct genetic linkage map of cowpea genome which can provide basic information towards genetic improvement of cowpea in breeding programmes.

## Acknowledgements

We are grateful for the support provided for this research by the Department of Plant and Environmental Biology, University of Ghana, Legon, Ghana Standard Authority and the Laboratory of the Government Chemist, United Kingdom.

## Supporting Information

**S1 Table :** Morphological traits and sources of cowpea accessions.

| ID | Plant type | Growth habit | Growth pattern | Twinning tendency | Plant pigment |
| --- | --- | --- | --- | --- | --- |
| R1 | F<sub>2</sub> (G X W) | Erect | Indeterminate | None | At the leaf base |
| R2 | F<sub>2</sub> (G X W) | Acute erect | Determinate | None | None |
| R3 | F<sub>2</sub> (G X W) | Semi-erect | Indeterminate | Pronounced | Slightly at the nodes |
| R4 | F<sub>2</sub> (G X W) | Climbing | Determinate | None | None |
| R5 | F<sub>2</sub> (G X W) | Climbing | Determinate | None | None |
| R6 | F<sub>2</sub> (G X W) | Erect | Determinate | None | None |
| R7 | F<sub>2</sub> (G X W) | Erect | Determinate | None | None |
| R8 | F<sub>2</sub> (G X W) | Climbing | Determinate | None | Slightly at the nodes |
| R9 | F<sub>2</sub> (G X W) | Erect | Determinate | None | None |
| R10 | F<sub>2</sub> (G X W) | Prostrate | Indeterminate | Pronounced | None |
| R11 | F<sub>2</sub> (G X W) | Climbing | Indeterminate | Pronounced | None |
| R12 | F<sub>2</sub> (G X W) | Prostrate | Indeterminate | Pronounced | Slightly at the nodes |
| R13 | F<sub>2</sub> (G X W) | Semi-prostrate | Indeterminate | Pronounced | Slightly at the nodes |
| R14 | F<sub>2</sub> (G X W) | Semi-prostrate | Indeterminate | Pronounced | Slightly at the nodes |
| R15 | F<sub>2</sub> (G X W) | Prostrate | Indeterminate | Pronounced | Slightly at the nodes |
| R16 | F<sub>2</sub> (G X W) | Semi-erect | Determinate | Pronounced | Slightly at the nodes |
| R17 | F<sub>2</sub> (G X W) | Semi-prostrate | Indeterminate | Pronounced | Slightly at the nodes |
| R18 | F<sub>2</sub> (G X W) | Acute erect | Determinate | None | None |
| R19 | F<sub>2</sub> (G X W) | Climbing | Determinate | None | None |
| R20 | F<sub>2</sub> (G X W) | Climbing | Determinate | None | None |
| R21 | F<sub>2</sub> (G X W) | Erect | Determinate | None | None |
| R22 | F<sub>2</sub> (G X W) | Erect | Determinate | None | None |
| R23 | F<sub>2</sub> (G X W) | Semi-erect | Determinate | None | None |
| R24 | F<sub>2</sub> (G X W) | Erect | Determinate | None | Slightly at the nodes |
| R25 | F<sub>2</sub> (G X W) | Semi-erect | Determinate | Pronounced | At the leaf base |
| R26 | F<sub>2</sub> (G X W) | Erect | Determinate | Pronounced | At the leaf base |
| R27 | F<sub>2</sub> (G X W) | Climbing | Determinate | Slightly pronounced | None |
| R28 | F<sub>2</sub> (G X W) | Semi-erect | Determinate | None | Slightly at the nodes |
| R29 | F<sub>2</sub> (G X W) | Erect | Determinate | Slightly pronounced | Intermediate |
| R30 | F<sub>2</sub> (G X W) | Prostrate | Indeterminate | Slightly pronounced | At the leaf base |
| R31 | F<sub>2</sub> (G X W) | Semi-prostrate | Indeterminate | Intermediate | Intermediate |
| R32 | F<sub>2</sub> (G X W) | Semi-prostrate | Indeterminate | Slightly pronounced | Intermediate |
| R33 | F<sub>2</sub> (G X W) | Semi-erect | Determinate | None | None |
| R34 | F<sub>2</sub> (G X W) | Climbing | Determinate | Slightly pronounced | Slightly at the nodes |
| R35 | F<sub>2</sub> (G X W) | Prostrate | Indeterminate | Pronounced | At the leaf base |
| R36 | F<sub>2</sub> (G X W) | Climbing | Determinate | None | None |
| R37 | F<sub>2</sub> (G X W) | Erect | Determinate | None | Intermediate |
| R38 | F<sub>2</sub> (G X W) | Semi-prostrate | Indeterminate | Intermediate | None |
| R39 | F<sub>2</sub> (G X W) | Erect | Determinate | None | None |
| R40 | F<sub>2</sub> (G X W) | Semi-erect | Determinate | Pronounced | Slightly at the nodes |
| R41 | F<sub>2</sub> (G X W) | Semi-prostrate | Indeterminate | Pronounced | Slightly at the nodes |
| R42 | F<sub>2</sub> (G X W) | Climbing | Determinate | Slightly pronounced | Slightly at the nodes |
| R43 | F<sub>2</sub> (G X W) | Climbing | Indeterminate | Intermediate | Slightly at the nodes |
| R44 | F<sub>2</sub> (G X W) | Climbing | Determinate | None | Slightly at the nodes |
| R45 | F<sub>2</sub> (G X W) | Erect | Determinate | None | None |
| R46 | GOLINGA | Prostrate | Determinate | Pronounced | Slightly at the nodes |
| R47 | WILD | Climbing | Indeterminate | Slightly pronounced | At the leaf base |

**S1 Table: Morphological traits and sources of cowpea accessions (Continued)**
| ID | Plant type | Leaf size | Leaf shape | Plant hairiness | Leaf colour |
| --- | --- | --- | --- | --- | --- |
| R1 | F<sub>2</sub> (G X W) | Big | Hastate | Glabrescent | Light green |
| R2 | F<sub>2</sub> (G X W) | Medium | Sub-globose | Pubescent | Dark green |
| R3 | F<sub>2</sub> (G X W) | Big | Globose | Short appressed hairs | Dark green |
| R4 | F<sub>2</sub> (G X W) | Medium | Sub-globose | Glabrescent | Light green |
| R5 | F<sub>2</sub> (G X W) | Medium | Sub-globose | Glabrescent | Light green |
| R6 | F<sub>2</sub> (G X W) | Medium | Sub-globose | Glabrescent | Light green |
| R7 | F<sub>2</sub> (G X W) | Medium | Sub-globose | Glabrescent | Light green |
| R8 | F<sub>2</sub> (G X W) | Medium | Sub-globose | Pubescent | Dark green |
| R9 | F<sub>2</sub> (G X W) | Medium | Sub-globose | Glabrescent | Light green |
| R10 | F<sub>2</sub> (G X W) | Big | Globose | Pubescent | Dark green |
| R11 | F<sub>2</sub> (G X W) | Big | Globose | Pubescent | Dark green |
| R12 | F<sub>2</sub> (G X W) | Big | Globose | Pubescent | Light green |
| R13 | F<sub>2</sub> (G X W) | Big | Sub-globose | Glabrescent | Dark green |
| R14 | F<sub>2</sub> (G X W) | Big | Globose | Pubescent | Light green |
| R15 | F<sub>2</sub> (G X W) | Big | Globose | Pubescent | Dark green |
| R16 | F<sub>2</sub> (G X W) | Big | Sub-hastate | Glabrescent | Dark green |
| R17 | F<sub>2</sub> (G X W) | Small | Sub-hastate | Glabrescent | Dark green |
| R18 | F<sub>2</sub> (G X W) | Medium | Sub-globose | Short appressed hairs | Light green |
| R19 | F<sub>2</sub>(G X W) | Medium | Sub-globose | Glabrescent | Dark green |
| R20 | F<sub>2</sub>(G X W) | Medium | Sub-globose | Glabrescent | Dark green |
| R21 | F<sub>2</sub>(G X W) | Small | Sub-globose | Glabrescent | Dark green |
| R22 | F<sub>2</sub>(G X W) | Medium | Sub-globose | Glabrescent | Dark green |
| R23 | F<sub>2</sub>(G X W) | Big | Globose | Pubescent | Dark green |
| R24 | F<sub>2</sub>(G X W) | Medium | Sub-globose | Glabrescent | Light green |
| R25 | F<sub>2</sub>(G X W) | Big | Hastate | Glabrescent | Dark green |
| R26 | F<sub>2</sub>(G X W) | Big | Sub-hastate | Glabrescent | Light green |
| R27 | F<sub>2</sub>(G X W) | Big | Globose | Glabrescent | Light green |
| R28 | F<sub>2</sub>(G X W) | Medium | Sub-globose | Short appressed hairs | Light green |
| R29 | F<sub>2</sub>(G X W) | Narrow | Hastate | Pubescent | Variegated |
| R30 | F<sub>2</sub>(G X W) | Narrow | Hastate | Short appressed hairs | Dark green |
| R31 | F<sub>2</sub>(G X W) | Narrow | Hastate | Pubescent | Light green |
| R32 | F<sub>2</sub>(G X W) | Medium | Sub-hastate | Glabrescent | Light green |
| R33 | F<sub>2</sub>(G X W) | Big | Globose | Short appressed hairs | Dark green |
| R34 | F<sub>2</sub>(G X W) | Medium | Sub-globose | Glabrescent | Light green |
| R35 | F<sub>2</sub>(G X W) | Narrow | Hastate | Pubescent | Dark green |
| R36 | F<sub>2</sub>(G X W) | Medium | Sub-globose | Short appressed hairs | Light green |
| R37 | F<sub>2</sub>(G X W) | Medium | Hastate | Pubescent | Dark green |
| R38 | F<sub>2</sub>(G X W) | Medium | Sub-globose | Pubescent | Dark green |
| R39 | F<sub>2</sub>(G X W) | Medium | Sub-hastate | Glabrescent | Light green |
| R40 | F<sub>2</sub>(G X W) | Medium | Sub-hastate | Glabrescent | Light green |
| R41 | F<sub>2</sub>(G X W) | Medium | Sub-hastate | Pubescent | Dark green |
| R42 | F<sub>2</sub>(G X W) | Medium | Sub-hastate | Short appressed hairs | Dark green |
| R43 | F<sub>2</sub>(G X W) | Medium | Sub-globose | Glabrescent | Light green |
| R44 | F<sub>2</sub>(G X W) | Medium | Sub-hastate | Pubescent | Light green |
| R45 | F<sub>2</sub>(G X W) | Medium | Sub-globose | Short appressed hairs | Light green |
| R46 | GOLINGA | Big | Globose | Glabrescent | Light green |
| R47 | WILD | Medium | Hastate | Pubescent | Dark green |

**S1 Table: Morphological traits and sources of cowpea accessions (Continued)**
| ID | Plant type | Leaf texture | Leaf markings | Petiole thickness | Raceme position |
| --- | --- | --- | --- | --- | --- |
| R1 | F<sub>2</sub> (G X W) | Cariaceous | Present | Thick | Uppermost canopy |
| R2 | F<sub>2</sub> (G X W) | Cariaceous | Absent | Intermediate | Throughtout canopy |
| R3 | F<sub>2</sub> (G X W) | Cariaceous | Present | Thick | Uppermost canopy |
| R4 | F<sub>2</sub> (G X W) | Cariaceous | Absent | Thick | Uppermost canopy |
| R5 | F<sub>2</sub> (G X W) | Cariaceous | Absent | Thin | Throughtout canopy |
| R6 | F<sub>2</sub> (G X W) | Cariaceous | Absent | Thin | Throughtout canopy |
| R7 | F<sub>2</sub> (G X W) | Intermediate | Present | Intermediate | Uppermost canopy |
| R8 | F<sub>2</sub> (G X W) | Cariaceous | Present | Thick | Uppermost canopy |
| R9 | F<sub>2</sub> (G X W) | Intermediate | Present | Thick | Uppermost canopy |
| R10 | F<sub>2</sub> (G X W) | Membranous | Absent | Thin | Throughtout canopy |
| R11 | F<sub>2</sub> (G X W) | Membranous | Present | Thick | Uppermost canopy |
| R12 | F<sub>2</sub> (G X W) | Cariaceous | Present | Thin | Throughtout canopy |
| R13 | F<sub>2</sub> (G X W) | Membranous | Absent | Thick | Uppermost canopy |
| R14 | F<sub>2</sub> (G X W) | Cariaceous | Present | Thin | Throughtout canopy |
| R15 | F<sub>2</sub> (G X W) | Membranous | Present | Thin | Throughtout canopy |
| R16 | F<sub>2</sub> (G X W) | Membranous | Present | Thin | Throughtout canopy |
| R17 | F<sub>2</sub> (G X W) | Cariaceous | Absent | Thick | Uppermost canopy |
| R18 | F<sub>2</sub> (G X W) | Cariaceous | Present | Thick | Uppermost canopy |
| R19 | F<sub>2</sub>(G X W) | Membranous | Absent | Thick | Uppermost canopy |
| R20 | F<sub>2</sub>(G X W) | Membranous | Absent | Thick | Uppermost canopy |
| R21 | F<sub>2</sub>(G X W) | Cariaceous | Absent | Thick | Uppermost canopy |
| R22 | F<sub>2</sub>(G X W) | Cariaceous | Absent | Thick | Uppermost canopy |
| R23 | F<sub>2</sub>(G X W) | Membranous | Present | Intermediate | Throughtout canopy |
| R24 | F<sub>2</sub>(G X W) | Cariaceous | Absent | Thin | Throughtout canopy |
| R25 | F<sub>2</sub>(G X W) | Membranous | Absent | Thick | Uppermost canopy |
| R26 | F<sub>2</sub>(G X W) | Membranous | Absent | Thick | Uppermost canopy |
| R27 | F<sub>2</sub>(G X W) | Cariaceous | Absent | Thick | Uppermost canopy |
| R28 | F<sub>2</sub>(G X W) | Cariaceous | Absent | Thin | Throughtout canopy |
| R29 | F<sub>2</sub>(G X W) | Cariaceous | Present | Thin | Throughtout canopy |
| R30 | F<sub>2</sub>(G X W) | Membranous | Present | Thick | Uppermost canopy |
| R31 | F<sub>2</sub>(G X W) | Cariaceous | Present | Thin | Throughtout canopy |
| R32 | F<sub>2</sub>(G X W) | Cariaceous | Absent | Thin | Throughtout canopy |
| R33 | F<sub>2</sub>(G X W) | Membranous | Present | Thin | Throughtout canopy |
| R34 | F<sub>2</sub>(G X W) | Cariaceous | Absent | Intermediate | Throughtout canopy |
| R35 | F<sub>2</sub>(G X W) | Membranous | Present | Thick | Uppermost canopy |
| R36 | F<sub>2</sub>(G X W) | Cariaceous | Absent | Thick | Uppermost canopy |
| R37 | F<sub>2</sub>(G X W) | Membranous | Present | Thick | Uppermost canopy |
| R38 | F<sub>2</sub>(G X W) | Intermediate | Present | Thin | Throughtout canopy |
| R39 | F<sub>2</sub>(G X W) | Membranous | Present | Thin | Throughtout canopy |
| R40 | F<sub>2</sub>(G X W) | Membranous | Present | Intermediate | Throughtout canopy |
| R41 | F<sub>2</sub>(G X W) | Membranous | Present | Thick | Uppermost canopy |
| R42 | F<sub>2</sub>(G X W) | Membranous | Present | Thin | Throughtout canopy |
| R43 | F<sub>2</sub>(G X W) | Cariaceous | Present | Thick | Uppermost canopy |
| R44 | F<sub>2</sub>(G X W) | Membranous | Absent | Thin | Throughtout canopy |
| R45 | F<sub>2</sub>(G X W) | Cariaceous | Present | Intermediate | Throughtout canopy |
| R46 | GOLINGA | Cariaceous | Present | Thin | Throughtout canopy |
| R47 | WILD | Membranous | Absent | Thick | Uppermost canopy |

